# Heme auxotrophy in abundant aquatic microbial lineages

**DOI:** 10.1101/2021.01.11.426183

**Authors:** Suhyun Kim, Ilnam Kang, Jin-Won Lee, Che-Ok Jeon, Stephen J. Giovannoni, Jang-Cheon Cho

## Abstract

Heme, a porphyrin ring complexed with iron, is a metalloprosthetic group of numerous proteins involved in diverse metabolic and respiratory processes across all domains of life, and is thus considered essential for respiring organisms^1,2^. Several microbial groups are known to lack the *de novo* heme biosynthetic pathway and therefore require exogenous heme from the environment^3^. These heme auxotroph groups are largely limited to pathogens^4,5^, symbionts^6,7^, or microorganisms living in nutrient-replete conditions^8^, whereas the complete absence of heme biosynthesis is extremely rare in free-living organisms^9^. Here, we show that the acI lineage, a predominant and ubiquitous free-living bacterial group in freshwater habitats, is auxotrophic for heme. We found that two recently cultivated acI isolates^10^ require exogenous heme for their growth. According to whole-genome analyses, all (*n*=20) isolated acI strains lacked essential enzymes necessary for heme biosynthesis, indicating that heme auxotrophy is a conserved trait in this lineage. Analyses of >24,000 representative genomes for species clusters of the Genome Taxonomy Database (GTDB) revealed that heme auxotrophy is widespread across abundant but not-yet-cultivated microbial groups, including *Patescibacteria*, *Marinisomatota* (SAR406), *Actinomarinales* (OM1), and marine group III *Euryarchaeota*. Our findings indicate that heme auxotrophy is a more common phenomenon than previously thought, and may lead to use of heme as a growth factor to increase the cultured microbial diversity.

---

We recently reported an approach for the robust cultivation of two acI strains^10^, thus establishing the first stable cultures of the actinobacterial acI lineage, one of the most abundant bacterioplankton groups in freshwater environments^11–13^. For long, efforts to cultivate the acI lineage using genome predictions and inferring ecophysiological traits were unsuccessful. Stable axenic cultivation of two acI strains was first achieved by supplying catalase as a critical growth supplement^10^, after which diverse sub-clades of the acI lineage^14^ were isolated with this approach. In this previous work^10^, the two strains, “*Candidatus* Planktophila rubra” IMCC25003 (hereinafter referred to as IMCC25003) and “*Candidatus* Planktophila aquatilis” IMCC26103 (hereinafter referred to as IMCC26103), grew robustly and reproducibly when bovine liver catalase was added to growth media. Since the growth was concomitant with decreases in H_2_O_2_ concentrations caused by the addition of catalase, it was suggested that the acI group might be sensitive to oxidative stress and the growth was due to lowered H_2_O_2_ concentrations. However, two experimental observations could not be easily reconciled with this interpretation. First, when H_2_O_2_ concentration was lowered by the addition of pyruvate, a chemical scavenger of H_2_O_2_, the enhancement of acI growth was marginal. Second, the H_2_O_2_ concentration that allowed the growth of the two acI strains in the catalase-supplemented laboratory experiment (~10 nM) was much lower than those measured in the lake from which the strains were isolated (~60 nM). Therefore, we conjectured that “catalase is essential for the growth of acI strains via mechanisms involving, but not restricted to, H_2_O_2_ decomposition”^10^.

While further investigating this unsolved discrepancy, we discovered that all genomes of the acI lineage were deficient in the heme biosynthetic pathway (**Extended Data Fig. 1**; discussed in more detail below), which was also reported in a recent study^15^. This finding led us to speculate that acI bacteria require heme, an essential cofactor of catalase, for their growth, rather than the H_2_O_2_ scavenging function of catalase. Here, based on genome predictions and experimental evidence, we report that the acI lineage is auxotrophic for heme. To our knowledge, these experiments with acI bacteria show for the first time that major and abundant lineages of free-living microorganisms inhabiting natural aquatic environments can be heme auxotrophs. We further demonstrate through comparative genomics that heme auxotrophy is found widely among many abundant prokaryotic groups.

All of twenty complete genome sequences from isolated acI strains were found to lack essential heme biosynthesis genes. The heme biosynthetic pathway is complex and consists of three parts^9^ (**Extended Data Fig. 1**). In the upper part, 5-amino-levulinic acid (ALA) is synthesized via two different modules known as C4 and C5. In the essential middle part, ALA is converted to uroporphyrinogen III (a common precursor of all heme compounds), via a three-step process common to all organisms, designated as “COMMON.” In the lower part, heme synthesis from uroporphyrinogen III is mediated by three different modules, each designated PPH (protoporphyrin-dependent), CPH (coproporphyrinogen-dependent), and SIRO (siroheme-dependent). Therefore, six variant pathways, combinations of one of the two upper modules with one of the three lower modules, are possible in theory for heme biosynthesis. All acI genomes lack any genes for the essential middle part of the pathway and also lack most genes for the upper and lower parts, which strongly indicates that these strains cannot synthesize heme and therefore require exogenous heme for survival. Hemes are iron-containing porphyrins that are present as cofactors in a diverse range of bacterial proteins, including respiratory cytochromes, non-respiratory-chain cytochromes, non-cytochrome hemoproteins, gas sensor proteins, peroxidases, and catalases^9^. Given that heme-containing cytochromes are essential for respiration, it was surprising to find that the acI lineage, which is comprised of free-living and obligately aerobic bacteria, does not possess complete sets of heme biosynthesis coding sequences.

As the first step to test heme auxotrophy of acI bacteria, we investigated the effect of hemin (the oxidized form of heme *b*) on the growth of the two acI strains, IMCC25003 and IMCC26103, belonging to the acI-A1 and acI-A4 tribes, respectively. IMCC25003 growth was dependent on hemin in a catalase-free medium. Growth was enhanced by hemin at concentrations as low as 10 pM, and enhancement was saturated at hemin concentrations of 1 nM or higher, with maximum cell densities (1.8–2.5 × 10^7^ cells mL^−1^) comparable to that obtained from bovine liver catalase addition (10 U mL^−1^) (**Fig. 1a**). Given that the heme concentration in 10 U mL^−1^ catalase is equivalent to ~50 nM, this amount of added catalase would be predicted to provide sufficient heme to support full IMCC25003 culture growth. On the other hand, hemin showed no effect on the growth of IMCC26103 (**Fig. 1b**).

**Fig. 1.**
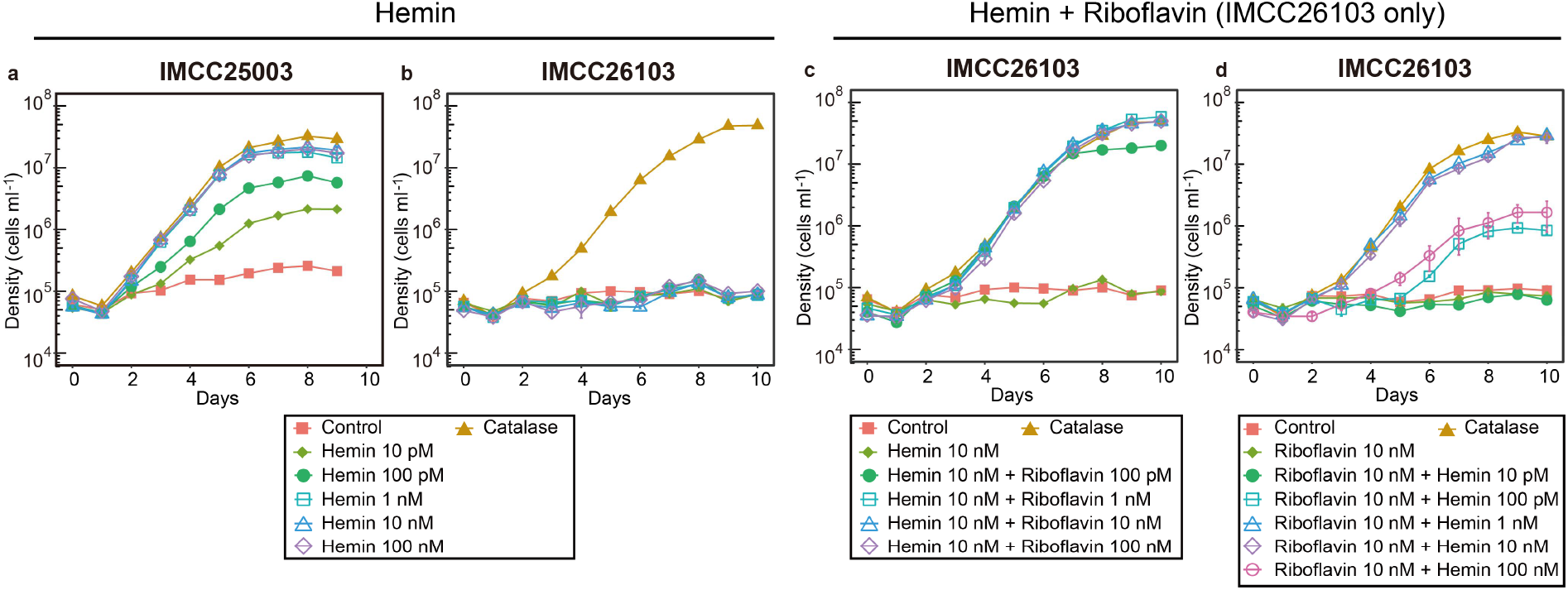
Hemin-dependent growth of the two acI strains. **a,b,** Effect of various concentrations (0–100 nM) of hemin on the growth of *’Ca.* P. rubra’ IMCC25003 (**a**) and *’Ca.* P. aquatilis’ IMCC26103 (**b**). **c**, Effect of various concentrations (0-100 nM) of riboflavin on the growth of IMCC26103 in the presence of hemin (10 nM). **d**, Effect of various concentrations (0-100 nM) of hemin on the growth of IMCC26103 in the presence of riboflavin (10 nM). The culture medium without hemin and riboflavin was designated as ‘Control’ and the medium with 10 U mL^−1^ of bovine liver catalase was designated as ‘Catalase’. All experiments were performed in triplicate. Error bars indicate the standard error. Note that error bars smaller than the size of the symbols are hidden.

Since hemin enhanced the growth of IMCC25003, but not IMCC26103, we hypothesized that IMCC26103 requires additional growth factor(s) that are present in catalase preparations and we predicted that, similar to IMCC25003, IMCC26103 would exhibit hemin-dependent growth if this additional factor was provided. A reinvestigation of genome-inferred metabolic potential^16^ identified riboflavin, a precursor of FMN and FAD, as the most likely additional growth factor required by IMCC26103 but not IMCC25003. The full gene set for *de novo* biosynthesis of riboflavin was identified only in IMCC25003. IMCC26103 instead had a gene for the riboflavin transporter PnuX, preceded by the FMN riboswitch (**Extended Data Fig. 2**). Further, the vitamin mixture used in the experiments did not contain riboflavin (**Extended Data Table 1**). Consistent with our genome-based predictions, IMCC26103 grew when riboflavin (≥100 pM) was provided together with hemin (10 nM), and its maximum cell densities in cultures supplemented with 1–100 nM of riboflavin reached to levels that were comparable to those of cultures supplied with catalase (**Fig. 1c**). Given that these results indicated that the catalase used in this study contained riboflavin, we directly measured the riboflavin concentration in the catalase solution. HPLC analysis of the catalase stock solution (10^5^ U mL^−1^) demonstrated that 10 U mL^−1^ of catalase contained approximately 690 pM of riboflavin (including FMN and FAD), which explains the growth enhancement of IMCC26103 in the catalase-supplemented cultures.

Upon demonstrating that IMCC26103 requires riboflavin for growth, the effect of hemin on the growth of the strain was reexamined in the presence of riboflavin (10 nM). IMCC26103 exhibited hemin-dependent growth (**Fig. 1d**), as shown for IMCC25003 (**Fig. 1a**), providing clear evidence of heme auxotrophy for both of the cultivated acI strains tested. Both acI strains grew well without catalase if the essential component of catalase, heme, and the impurity in the catalase stock, riboflavin, were supplied. It is worth noting that the relationships between growth and hemin concentration were different for the two strains. For instance, a hemin supply between 1 and 10 nM maximized the growth of both strains; however, a higher hemin concentration (≥100 pM) was required to grow IMCC26103 (in contrast to 10 pM for IMCC25003), and growth was reduced at 100 nM with IMCC26103, but not IMCC25003 **(Fig. 1a,d)**. The requirement of higher hemin concentration by IMCC26103 might be ascribed to a difference in putative heme transporters^17^. Upon searching for heme uptake systems in the acI genomes, gene clusters similar to the *dppABCDF* system^18,19^ were found in both strains, but a gene cluster similar to *hemTUV*^20,21^ was found only in IMCC25003, suggesting that acI bacteria may have a varied repertoire of heme uptake systems that result in differences in the minimal heme concentrations required for growth. We speculate that the reduced growth enhancement of IMCC26103 at higher hemin concentrations (100 nM) may be due to sensitivity to oxidative stress caused by excess heme (e.g., via the Fenton reaction^22,23^), as IMCC26103 lacks KatG.

Concentrations of dissolved iron–porphyrin-like complexes (up to 11.5 nM^24^) and heme *b* (47.3–70.8 pM^25^) in natural waters are similar to or higher than 10–100 pM, the minimum heme concentrations that supported the growth of acI strains, and therefore it seems likely that acI bacterioplankton survive in their habitats through uptake of exogenous heme. In natural systems, heme could originate from many sources, e.g. hemoproteins such as cytochromes, liberated from co-occurring organisms by diverse mechanisms, including viral lysis^26,27^. Additional measurements of dissolved heme concentration in aquatic habitats and the characterization of heme uptake systems of the acI lineage are required to better understand the effect of variable heme concentrations on acI populations.

With the establishment of robust growth conditions for acI strains by supplying heme and riboflavin, which do not degrade H_2_O_2_ (unlike catalase), we were able to reinvestigate the extreme growth sensitivity of the acI strains to H_2_O_2_ reported in our previous study. First, we used pyruvate (100 nM–50 μM) to reduce H_2_O_2_ concentrations in our basal medium (filtered and autoclaved lake water; ~600 nM of H_2_O_2_), as pyruvate is a H_2_O_2_-scavenging chemical and is not utilized by the acI strains as a carbon source^10^. IMCC25003 grew in all conditions, with a slight inhibition dependent on H_2_O_2_ concentration (**Fig. 2a**). Even at H_2_O_2_ concentrations above 200 nM throughout the incubation period (pyruvate: 0 or 100 nM), maximum cell densities reached ~6 × 10^6^ cells mL^−1^ (**Fig. 2a,e**). On the other hand, IMCC26103 exhibited growth only when ≥10 μM pyruvate was added, reducing the H_2_O_2_ concentration far below 200 nM, thus indicating that IMCC26103 is more sensitive to H_2_O_2_ than IMCC25003 (**Fig. 2b,f**). To assess the growth of the acI strains at higher H_2_O_2_ concentrations, the two strains were cultured in media supplemented with H_2_O_2_ (100 nM–10 μM). Although inhibited, IMCC25003 grew at initial H_2_O_2_ concentrations of nearly 1 μM, and complete inhibition was observed only when H_2_O_2_ remained above 5 μM (**Fig. 2c,g**). As expected, IMCC26103 failed to grow in any of the H_2_O_2_-supplemented conditions (**Fig. 2d,h**). These observations using culture media supplemented with heme, riboflavin, and varying H_2_O_2_ concentrations demonstrate that the acI strains investigated herein are not as sensitive to H_2_O_2_ as previously thought. We postulate that these differences in H_2_O_2_ sensitivity between the two acI strains are due to the presence of *katG* (a gene for bifunctional catalase-peroxidase) in IMCC25003, but not in IMCC26103. The KatG protein of IMCC25003 displayed catalase activity in our previous work^10^, and therefore its presence in IMCC25003 likely explains why this strain can defend itself against H_2_O_2_ stress better than IMCC26103. This difference could lead to niche partitioning among the acI strains, as large spatiotemporal variation in H_2_O_2_ concentrations has been reported in freshwater habitats^28^.

**Fig. 2.**
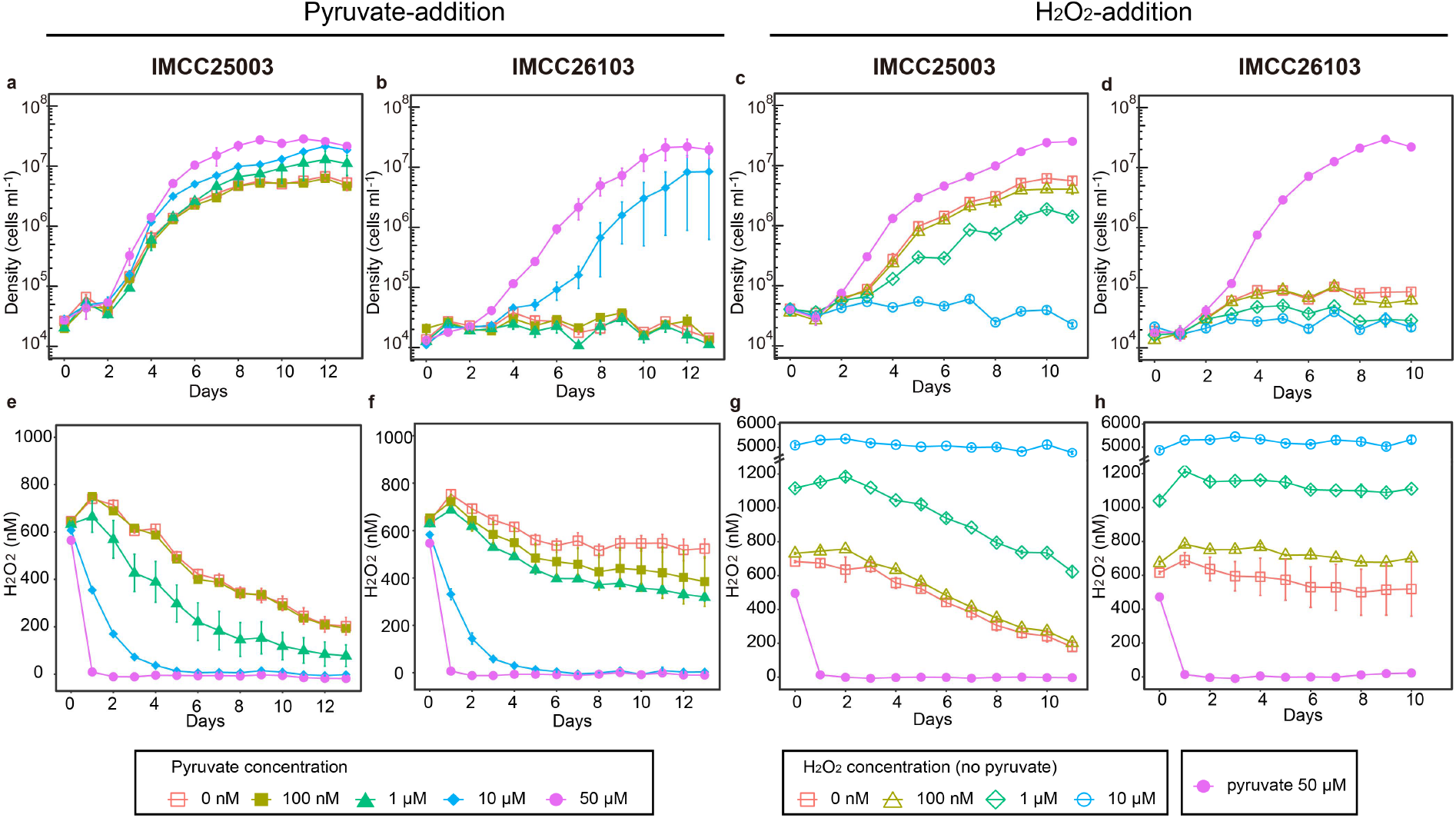
Effects of H_2_O_2_ on the growth of acI strains. Various concentrations of pyruvate (0–50 μM; **a, b, e, f**) and H_2_O_2_ (0–10 μM; **c, d, g, h**) were added to the basal media containing hemin (10 nM) and riboflavin (1 nM) to adjust the hydrogen peroxide concentration of the media. **a–d**, Growth of IMCC25003 (**a,c**) and IMCC26103 (**b,d**). **e–h**, Changes in H_2_O_2_ concentration during cultivation of IMCC25003 (**e,g**) and IMCC26103 (**f,h**). All experiments were performed in triplicate. Error bars indicate the standard error. Note that error bars shorter than the size of the symbols are hidden.

To gain insights into the mechanisms by which the acI lineage lost the heme biosynthetic pathway, we analyzed the distribution of the pathway in the genomes of *Nanopelagicales*. In the GTDB (Genome Taxonomy Database)^29^, the acI lineage is classified as *Nanopelagicaceae* within the order *Nanopelagicales* constituting several families. This analysis was performed by taking the maximum value of the completeness of six variants of heme biosynthetic pathways found in each representative genome for species clusters within *Nanopelagicales* available in the GTDB (release 89) (**Extended Data Fig. 1, Extended Data Table 1**). The pathway was almost completely deficient in all freshwater *Nanopelagicaceae* genomes, including single-amplified genomes (SAGs) and metagenome-assembled genomes (MAGs), whereas only one MAG from permafrost soil (palsa_747) contained genes encoding early steps in the pathway (**Fig. 3a**). Other *Nanopelagicales* genomes not affiliated with the acI lineage were more complete than the acI genomes. Particularly, three MAGs from the ocean (NP52 and NAT138) or wastewater (UBA10799) had nearly complete pathways (**Fig. 3a**). Furthermore, given that most of the *Actinobacteria* genomes exhibit the complete heme biosynthetic pathway (**Extended Data Fig. 3**), it seemed plausible that the acI lineage had lost the heme biosynthetic pathway while adapting to freshwater environments through genome streamlining. Discarding the heme biosynthetic pathway could, in principle, be a risky strategy for free-living aerobes because many essential biochemical reactions rely on hemoproteins (e.g., oxidative phosphorylation). Therefore, cells lacking the heme biosynthetic pathway would be destined to depend on exogenous heme, the supply of which could be unstable. However, the few studies that have measured free dissolved heme or heme-like molecules in aquatic habitats have detected concentrations that would be sufficient to support the growth of the acI strains studied herein. Fitness benefits from abandoning heme biosynthesis likely favor auxotrophic dependency. Aside from saving the resources that would be needed to maintain and express the heme biosynthetic machinery, auxotrophic cells might also be avoiding costs associated with heme toxicity. Excess heme is toxic to living organisms via a multitude of mechanisms (e.g., the Fenton reaction caused by iron liberated from heme)^22,23^. Ostensibly to counter this toxicity, cells finely regulate heme biosynthesis and have strategies for heme detoxification, such as efflux systems^30,31^. The metabolic burdens from regulation and detoxification might be partially diminished by abandoning heme biosynthesis.

**Fig. 3.**
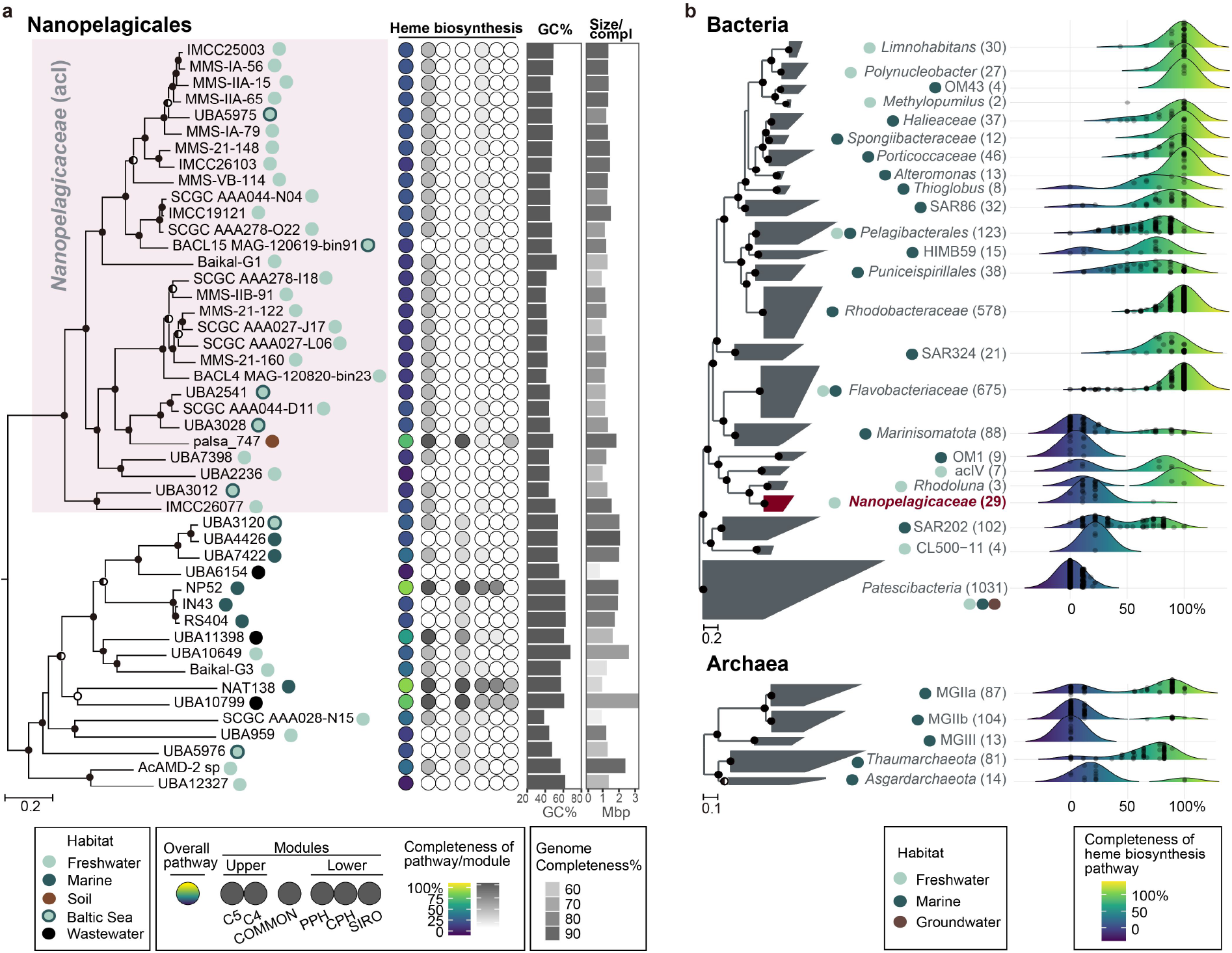
Heme biosynthetic pathway completeness of *Nanopelagicales* and major aquatic prokaryotic groups. **a,** Heme biosynthetic pathway completeness of representative genomes for species clusters belonging to *Nanopelagicales*. A phylogenomic tree of *Nanopelagicales* constructed using a concatenated alignment of conserved marker proteins is shown on the left. The circles on the right side of genome names indicate isolation sources (habitats). The *Nanopelagicaceae* (acI) genomes are shaded in pink. *Rhodoluna lacicola* MWH-Ta8 (not shown here) was set as an outgroup. The middle column illustrates the overall (leftmost circle) and per-module (the remaining six circles) completeness of the heme biosynthetic pathway among the *Nanopelagicales* genomes. Refer to Extended Data Fig. 1 for the heme biosynthetic pathway completeness calculations. The circle colors indicate the pathway completeness according to the color gradients in the bottom legend. The GC contents and genome sizes of the genomes are shown with bar graphs on the right. Genome completeness is indicated by the darkness of the genome size bars, as indicated by the scale at the bottom. **b**, Distribution of the completeness of the heme biosynthetic pathway in major aquatic prokaryotic groups. Phylogenomic trees of representative genomes for species clusters affiliated with major aquatic bacterial and archaeal groups (Extended Data Table 2) are shown on the left. The generally recognized habitats and the number of genomes of the groups are indicated at the left and right (within parentheses) of the group names, respectively. The *Nanopelagicaceae* family (acI lineage) is indicated in red. The phylum *Patescibacteria* and phyla *Thaumarchaeota* and *Asgardarchaeota* were set as the outgroups in the bacterial and archaeal trees, respectively. On the right, ridgeline density plots were used to illustrate the distribution of the completeness of heme biosynthetic pathways in the prokaryotic groups. The colors under the ridgelines indicate the completeness following a color gradient shown at the bottom. The points in the plots indicate the completeness of each genome of the groups. For the trees in this figure, bootstrap values (a; from 100 resamplings) and local support values (b; from 1,000 resamplings) are shown at the nodes as filled circles (≥90%), half-filled circles (≥70%), and empty circles (≥50%).

Having confirmed heme auxotrophy in the acI lineage, we expanded the scope of our investigation to assess genomic evidence for this trait among all bacteria and archaea. This analysis targeted a total of 24,706 bacterial and archaeal genomes representing species clusters of the GTDB database (**Supplementary Table 1**). We found that many phyla consisted exclusively of members that lacked heme biosynthesis genes (e.g., *Dependentiae*, *Patescibacteria*, and DPANN containing *Nanoarchaeota*) or contained substantial proportions of predicted heme auxotrophs [e.g., *Marinisomatota* (SAR406), *Chloroflexota*, and *Asgardarchaeota*] (**Extended Data Figs. 3, 4**). Many of the phyla in which heme biosynthetic pathway gaps were frequent consisted of not-yet or rarely cultured organisms, the so-called microbial dark matter^32^, suggesting that heme auxotrophy is a plausible explanation for the challenges of culturing the uncultured microbial majority. Alternatively, considering the enormous metabolic diversity of prokaryotes, unknown heme biosynthesis pathways are a possibility, and several of the groups predicted to be heme-auxotrophs (e.g., *Patescibacteria* and DPANN) are inferred from their genomes to lack respiration and rely strictly on fermentative metabolism^33,34^; these cells could conceivably replicate with little to no exogenous heme.

The apparent heme auxotrophy in several abundant planktonic clades, such as SAR406 and Marine Group II archaea (MG-II), led us to analyze aquatic microorganisms in more detail (**Fig. 3b**). A total of 3,233 representative genomes for species clusters of 29 major phylogenetic groups residing in marine, freshwater, and groundwater environments were analyzed. The selected groups and their GTDB taxonomy are listed in **Extended Data Table 2**. Evidence of deficiencies in heme biosynthetic pathways was widespread, although varying in degree, in many not-yet-cultivated groups such as SAR406, *Actinomarinales* (OM1), CL500-11, *Patescibacteria*, MG-IIb, MG-III, and *Asgardarchaeota* (**Fig. 3b**). Inspection of phylogenomic trees indicated that heme-auxotrophy is relegated to specific sub-clades within each of these groups (**Extended Data Fig. 5-11**), which indicates that i) this evidence for heme auxotrophy is unlikely to be an artifact caused by the incompleteness of SAGs and MAGs, and ii) the loss of heme biosynthetic pathways likely occurred independently many times during evolutionary adaptation to specific environmental niches.

Our findings confirmed heme-auxotrophy in the predominant freshwater microbe acI and uncovered genomic evidence indicating that heme-auxotrophy has evolved repeatedly among microorganisms from many environments, particularly among cosmopolitan aquatic plankton. The previously reported effect of catalase on the growth of acI^10^ was found to be due to heme, an essential cofactor of catalase, rather than the enzymatic activity of catalase. Our findings add to a growing list of examples in which cells dispense with expensive metabolic pathways in favor of increased reliance on compounds that are fundamentally important to their metabolism but are synthesized by members of surrounding microbial communities. Two important themes in microbiome science converge in the findings we report: the connectivity of microbiomes and the challenge of culturing a broader range of the uncultured microbial majority. The motivations for advancing the knowledge on these topics are the need to expand the range of cell types that can be studied using experimental microbial cell biology approaches, and also the need to understand the rules that govern interactions among the members of microbial communities^35^. The discovery of widespread heme reliance in microbial ecosystems will likely stimulate future investigations of heme concentrations across biomes and the influence of heme traffic between cells on the structuring of microbial communities.

## Methods

### Cultivation of the acI strains

The basal culture medium used in this study was prepared by adding carbon compounds, nitrogen and phosphorus sources, an amino acid mixture, a vitamin mixture, and trace metals (**Extended Data Table 1**)into 0.2 μm-filtered and autoclaved lake water^10^. The lake water used to prepare the medium was collected from the surface (1 m) of Lake Soyang in January 2020. Strains IMCC25003 and IMCC26103 were maintained aerobically in the basal medium supplemented with catalase (10 U mL^−1^) at 20°C. A catalase stock solution (10^5^ U mL^−1^ in 10 mM PBS, pH 7.4) was prepared using bovine liver catalase (C9322, Sigma-Aldrich). For the growth experiments, stock solutions of hemin (5 mM in 5 mM NaOH; 51280, Sigma-Aldrich) and riboflavin (500 μM in Milli-Q water; R9504, Sigma-Aldrich) were prepared and sterilized with filtration through a 0.1 μm pore-size membrane filter. A H_2_O_2_ solution (H1009, Sigma-Aldrich) was diluted at a 1:10,000 ratio with sterile water. A pyruvic acid stock solution (50 mM) was also prepared and sterilized with a 0.1 μm pore-size membrane filter prior to use. All culture experiments were performed at 20°C in the dark using acid-washed polycarbonate flasks. Actively growing cells in the late exponential phase were used as inocula. Cell counts were obtained using a flow cytometer (Guava easyCyte Plus, Millipore) after staining with SYBR Green I (final concentration 5X; Invitrogen) for 1 hr.

### Determination of riboflavin concentration

HPLC analysis was conducted to quantify the amount of riboflavin in the catalase stock solution. A total of 5 mL of catalase stock solution was diluted to 50 mL with Milli-Q water and heated at 80°C in a water bath for 20 min, then analyzed with an HPLC system coupled with a fluorescence detector (Agilent Co.) and a CAPCELL PAK C18 MG (250×;4.6 mm, 5 μm, Shiseido) column. The detector was set to 445 nm excitation and 530 nm emission wavelengths. The mobile phase was prepared with 10 mM NaH2PO4 (pH 5.5) and methanol (76:24, v/v), and the analysis was conducted with 10 μl of injection volume at a 0.9 mL min^−1^ flow rate for 30 min. The amount of riboflavin, FMN, and FAD in the sample was calculated based on the peak area of the chromatogram, using standard curves with a high R^2^ value (0.999) obtained when FAD, FMN, and riboflavin standards were separated under the same conditions. The riboflavin, FMN, and FAD used for the determination of standard curves were purchased from Sigma-Aldrich Co.

### Measurement of hydrogen peroxide concentration

Hydrogen peroxide concentration was analyzed using the Amplex Red hydrogen peroxide/peroxidase assay kit (A22188, Invitrogen) according to the manufacturer’s instructions. For the assay, 100 μL of Amplex Red working solution (prepared according to the manufacturer’s instructions) was added to each assay tube containing 100 μL of sample. After incubation for 5 min at room temperature in the dark, the fluorescence of resorufin (the oxidized form of the Amplex Red reagent) was measured at excitation and emission wavelengths of 470 nm and 665–720 nm, respectively, using a Qubit 3 fluorometer (Thermo Fisher Scientific). To quantify H_2_O_2_ concentrations, a standard curve (0.01–25 μM of H_2_O_2_) was generated by diluting 3.6% (v/v) of H_2_O_2_ solution with sodium phosphate buffer supplied with the kit.

### Analysis on the completeness of the heme biosynthetic pathway in prokaryotic genomes

A total of 24,706 representative genome sequences for species clusters in the GTDB (Release 89; 23,458 Bacteria and 1,248 Archaea) were downloaded from the GTDB server^29^. Additionally, the genome sequences of the 20 acI strains isolated to date were downloaded from GenBank. Annotation of the genome sequences was performed using Prokka (v1.14.6), with no rRNA, no tRNA, and no annotation options^36^. Predicted protein sequences were searched against the KofamKOALA database (accessed on Feb 19, 2020) using KofamScan 1.2.0^37^. To reduce run time, the KofamScan was executed using a custom-built hal file containing only 25 KEGG Ontology (KO) IDs involved in heme biosynthetic pathways (**Extended Data Fig. 1**), with the ‘-f mapper’ option to retain only the hits above thresholds specific to KO IDs^38^. The 25 KO IDs were curated based on a KEGG pathway map titled “Porphyrin and chlorophyll metabolism (ko00860)”, KEGG modules M00121 (heme biosynthesis, plants and bacteria, glutamate => heme), M00868 (heme biosynthesis, animals and fungi, glycine => heme), M00846 (siroheme biosynthesis, glutamyl-tRNA => siroheme), and M00847 (heme biosynthesis, archaea, siroheme => heme), in addition to several previous studies^9,39^. The KofamScan output files were then merged for the next step.

To calculate the completeness of the heme biosynthetic pathway using the KofamScan output, we modified “KEGG_decoder.py” script^40^ using custom-built definitions for the 6 modules (C5, C4, COMMON, PPH, CPH, and SIRO) and the 6 variants (C5_PPH, C5_CPH, C5_SIRO, C4_PPH, C4_CPH, and C4_SIRO) of the pathway (**Extended Data Fig. 1**). These pathway definitions are based on the curated pathways and the 25 associated KOs presented in **Extended Data Fig. 1**. The modified Python script (available at https://github.com/SuhyunInha/KEGGdecoder_Heme) was executed with the merged KofamScan output file as an input, which produced completeness values for each of the six modules and six variants for all genomes. For each genome, the maximum completeness value among the six variants was taken as the overall heme biosynthetic pathway completeness.

### Reconstruction of phylogenomic trees and visualization of the heme biosynthetic pathway completeness

To visualize the distribution of the heme biosynthetic pathway completeness in prokaryotic genomes, phylogenomic trees were constructed using the trimmed and concatenated multiple sequence alignment of 120 bacterial and 122 archaeal marker proteins downloaded from the GTDB (Release 89). Two phylum-level trees (**Extended Data Fig. 3,4**) were inferred by FastTree2 with the default options^41^ using all the representative genomes for species clusters. Phyla containing 10 or fewer genomes were not included in the tree building for visualization. The tree in **Fig. 3b** was also built using FastTree2, using the genomes affiliated with the phylogenetic groups listed in **Extended Data Table 2**. All other trees were built using RAxML 8.2.7 with the PROTGAMMAAUTO option^42^. Genome statistics including completeness and contamination calculated by CheckM, size, and GC content were downloaded from the GTDB. Visualization of pathway completeness (modules and variants) was performed using the tidyverse^43^, ggridges^44^, and viridis packages in R environment (version 4.0.2)^45^.

## Acknowledgments

This study was supported by the Mid-Career Research Program (NRF-2019-R1A2B5B02070538 to J-CC) and Science Research Center grant of the NRF (No. NRF-2018R1A5A1025077 to J-CC, J-WL, C-OK) through the National Research Foundation (NRF) funded by the Ministry of Sciences and ICT, and by the Basic Science Research Program funded by the Ministry of Education, Republic of Korea (NRF-2020R1A6A3A01100626 to S.K. and NRF-2019R1I1A1A01063401NRF to I.K.).

## Author contributions

S.K. performed the wet experiments. S.K and I.K performed all bioinformatic data analyses. I.K. and J.-C.-C designed and directed the study. S.K., I.K., J.-W.L, C.-O.J, S.J.G, and J.-C.C. analyzed the data and wrote the manuscript.

