## Extended figures and tables for "Heme auxotrophy in abundant aquatic microbial lineages"

**a**

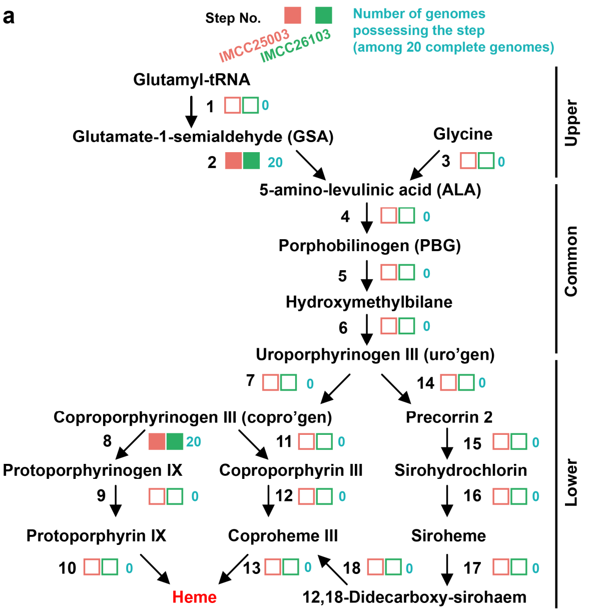


**b**

List of enzymes

| Step | Enzyme | KO ("\|" → OR) |
|---|---|---|
| 1 | Glu-tRNA reductase (HemA) | K02492 |
| 2 | GSA-2,1-aminomutase (HemL) | K01845 |
| 3 | 5-aminolevulinate synthase | K00643 |
| 4 | ALA dehydratase (HemB) | K01698 |
| 5 | PBG deaminase (HemC) | K01749 |
| 6 | uro'gen synthase (HemD) | K01719 \| K13542 \| K13543 |
| 7 | uro'gen decarboxylase (HemE) | K01599 |
| 8 | copro'gen oxidase (HemF/N) | K00228 \| K02495 |
| 9 | protoporphyrinogen oxidase (HemG/J/Y) | K00230 \| K00231 \| K08973 |
| 10 | protoporphyrin ferrochelatase (HemH) | K01772 |
| 11 | protoporphyrinogen oxidase (HemY) | K00231 |
| 12 | ferrochelatase (HemH) | K01772 |
| 13 | iron-coproporphyrin decarboxylase (HemQ) | K00435 \| K22227 |
| 14 | uroporphyrin -III C-methyltransferase (MET1, cysG, cobA, hemX, cobA-hemD, hemDX) | K00589 \| K02302 \| K02303 \| K02496 \| K13542 \| K13543 |
| 15 | precorrin-2 dehydrogenase (cysG, MET8) | K02302 \| K02304 |
| 16 | sirohydrochlorin ferrochelatase (cysG, MET8, sirB) | K02302 \| K02304 \| K03794 |
| 17 | siroheme decarboxylase (ahbAB) | K22225 |
| 18 | Fe-coproporphyrin III synthase (ahbC) | K22226 |

**c** Definition of six modules involved in heme biosynthesis in Prokaryote

| Parts | Modules | Step | Phylogenetic distribution |
|---|---|---|---|
| Upper | C5 | 1+2 | Plants, archaea, most bacteria |
| | C4 | 3 | Metaxoans, fungi, Alphaproteobacteria |
| Common | COMMON | 4+5+6 | Universal |
| Lower | PPH (Protoporphyrin-dependent) | 7+8+9+10 | Gram-negative bacteria |
| | CPH (Coproporphyrin-dependent) | 7+11+12+13 | Gram-positive bacteria |
| | SIRO (Siroheme-dependent) | 14+15+16+17+18 | Some Archaea and sulfate-reducing bacteria |

Six possible pathway variants for heme biosynthesis in Prokaryote

| Pathway variant (PV) | Upper | Common | Lower |
|---|---|---|---|
| C5_PPH | 1 + 2 (C5) | 4 + 5 + 6 | 7 + 8 + 9 + 10 (PPH) |
| C5_CPH | 1 + 2 (C5) | 4 + 5 + 6 | 7 + 11 + 12 + 13 (CPH) |
| C5_Siro | 1 + 2 (C5) | 4 + 5 + 6 | 14 + 15 + 16 + 17 + 18 + 13 (SIRO) |
| C4_PPH | 3 (C4) | 4 + 5 + 6 | 7 + 8 + 9 + 10 (PPH) |
| C4_CPH | 3 (C4) | 4 + 5 + 6 | 7 + 11 + 12 + 13 (CPH) |
| C4_SIRO | 3 (C4) | 4 + 5 + 6 | 14 + 15 + 16 + 17 + 18 + 13 (SIRO) |

**d**

$$\textbf{Module completeness (\%)} = \frac{\text{Number of steps found in genome}}{\text{Number of total steps (from 1 to 5)}} \times 100$$

$$\textbf{Completeness of pathway variants (\%)} = \frac{\text{Number of steps found in genome}}{\text{Number of total steps (from 8 to 11)}} \times 100$$

**Overall completeness of heme biosynthetic pathway** = Maximum of the completeness of 6 pathway variants

**Extended Data Fig. 1** Overview of heme biosynthetic pathways in prokaryotes and the acI lineage. **a**, Diagram showing the heme biosynthetic pathway steps. The presence (filled) or absence (empty) of each step in the two acI genomes are indicated by colored boxes (orange, IMCC25003; green, IMCC26103). The numbers of acI genomes (among 20) possessing each step are written in blue. **b**, List of enzyme names and corresponding KO IDs of the steps defined in the diagram. **c**, Definition of modules and heme biosynthetic pathway variants used in this study. **d**, Formulae for the calculation of completeness of modules and pathway variants. The maximum value among the completeness of six pathway variants of each genome was taken as the overall completeness of the genome.

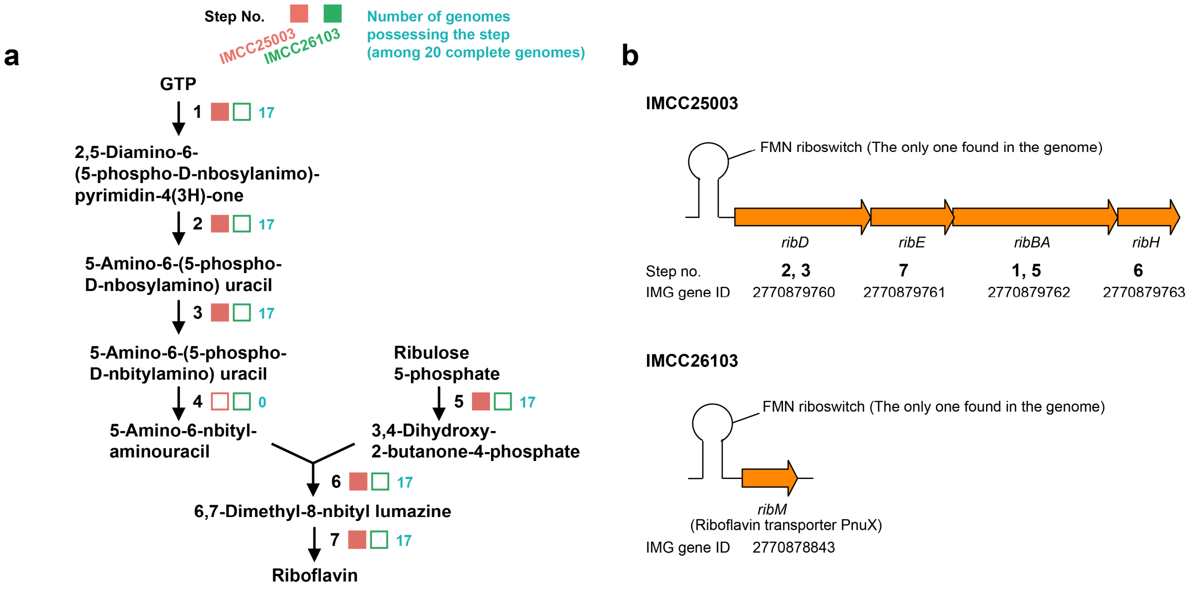


**Extended Data Fig. 2** Riboflavin biosynthetic pathway in the acI genomes. **a**, The presence (filled) and absence (empty) of each step in the two acI genomes are indicated with colored boxes (orange, IMCC25003; green, IMCC26103). The numbers of the acI genomes (among 20) possessing each step were written in blue. Note that the putative enzymes mediating step four, the only missing step in IMCC25003, have only recently been identified[1,2], and therefore many genomes of riboflavin-prototrophic microorganisms are still predicted to lack this step according to KEGG-based annotation. **b**, The genes for riboflavin biosynthesis (IMCC25003) and transport (IMCC26103) are located just downstream of the only FMN riboswitch predicted in each genome.

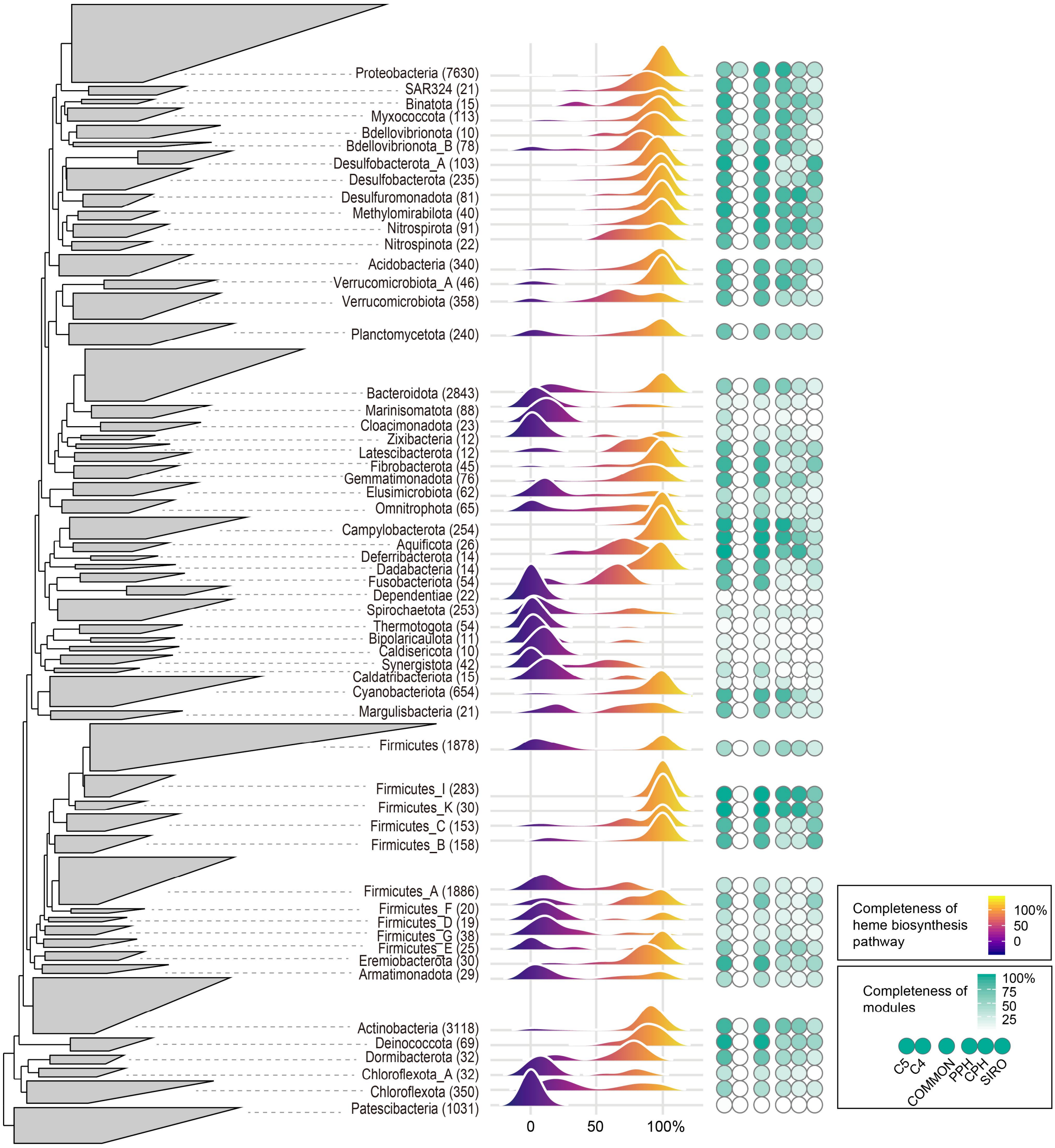


**Extended Data Fig. 3** Distribution of the heme biosynthetic pathway completeness in diverse bacterial phyla. **left**, An approximate maximum-likelihood tree of representative genomes for bacterial species clusters constructed by FastTree2 using a concatenated alignment of 120 conserved marker proteins. The genomes were grouped by phyla, and the phylum *Patescibacteria* was set as an outgroup. Phyla with less than 10 genomes were excluded from tree building. **middle,** Ridgeline density plots showing the distribution of the heme biosynthetic pathway completeness in bacterial phyla. The colors under the ridgelines indicate the pathway completeness according to the color gradient on the right. **right,** Average completeness of six pathway modules of bacterial phyla. The color intensity indicates the completeness according to the color gradient on the right. Refer to

Extended Data Fig. 1 for more details on the pathway modules.

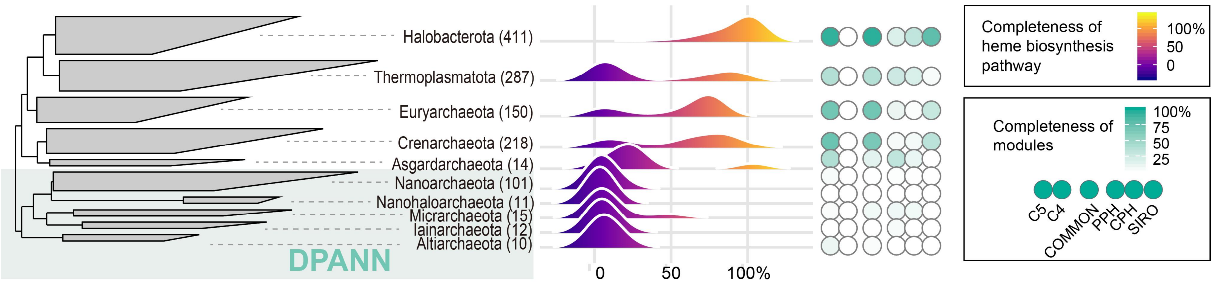


**Extended Data Fig. 4** Distribution of the heme biosynthetic pathway completeness in diverse archaeal phyla. **left**, An approximate maximum-likelihood tree of representative genomes for archaeal species clusters constructed by FastTree2 using a concatenated alignment of 122 conserved marker proteins. The genomes were grouped by phyla, and the superphylum DPANN (from *Nanoarchaeota* to *Altiarchaeota*) was set as an outgroup. Phyla with less than 10 genomes were excluded from tree building. **middle,** Ridgeline density plots showing the distribution of the heme biosynthetic pathway completeness in archaeal phyla. The colors under the ridgelines indicate pathway completeness according to the color gradient on the right. **right,** Average completeness of six pathway modules of archaeal phyla. The color intensity indicates the pathway completeness according to the color gradient on the right. Refer to Extended Data Fig. 1 for more details on the pathway modules.

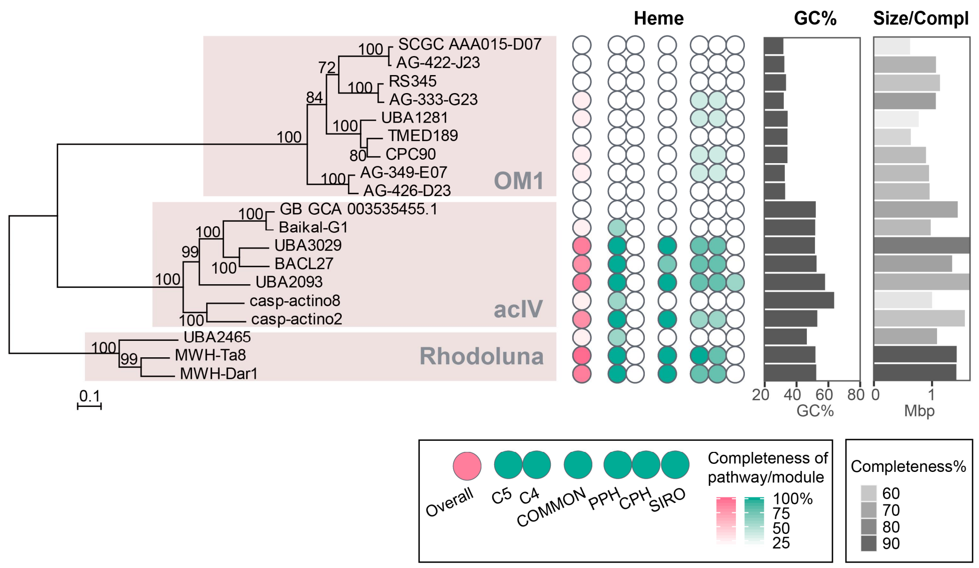


**Extended Data Fig. 5** Heme biosynthetic pathway completeness of representative genomes for species clusters belonging to OM1 (*Candidatus* Actinomarinales; o__TMED189 in GTDB), acIV, and the genus *Rhodoluna* of *Actinobacteriota*. **left**, Maximum-likelihood tree constructed by RAxML using a concatenated alignment of conserved marker proteins. The genus *Rhodoluna* was set as an outgroup. **middle,** Overall heme biosynthetic pathway completeness (indicated in pink) and the completeness of six pathway modules (indicated in green) within the genomes. The color intensity indicates the pathway completeness according to the color scales in the bottom legend. **right,** The GC contents and genome sizes are illustrated with bar graphs. Genome completeness is indicated as the darkness of the genome size bars according to the gray scale at the bottom. The data were downloaded from the GTDB (Release 89).

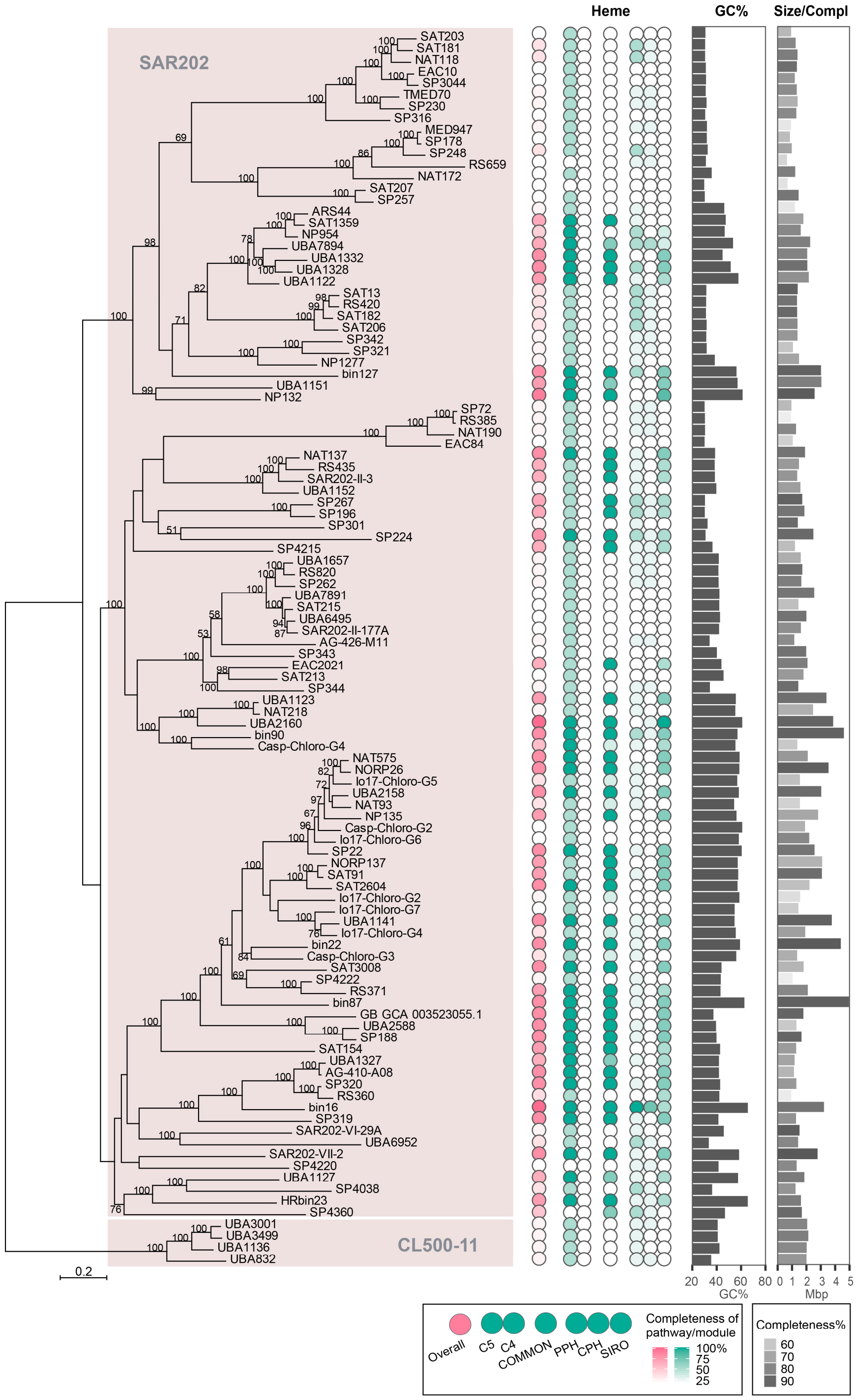


**Extended Data Fig. 6** Heme biosynthetic pathway completeness of representative genomes for species clusters belonging to the SAR202 group of *Chloroflexota*. The CL500-11 group was set as an outgroup in the tree on the left. Refer to the Extended Data Fig. 5 legend for a detailed explanation.

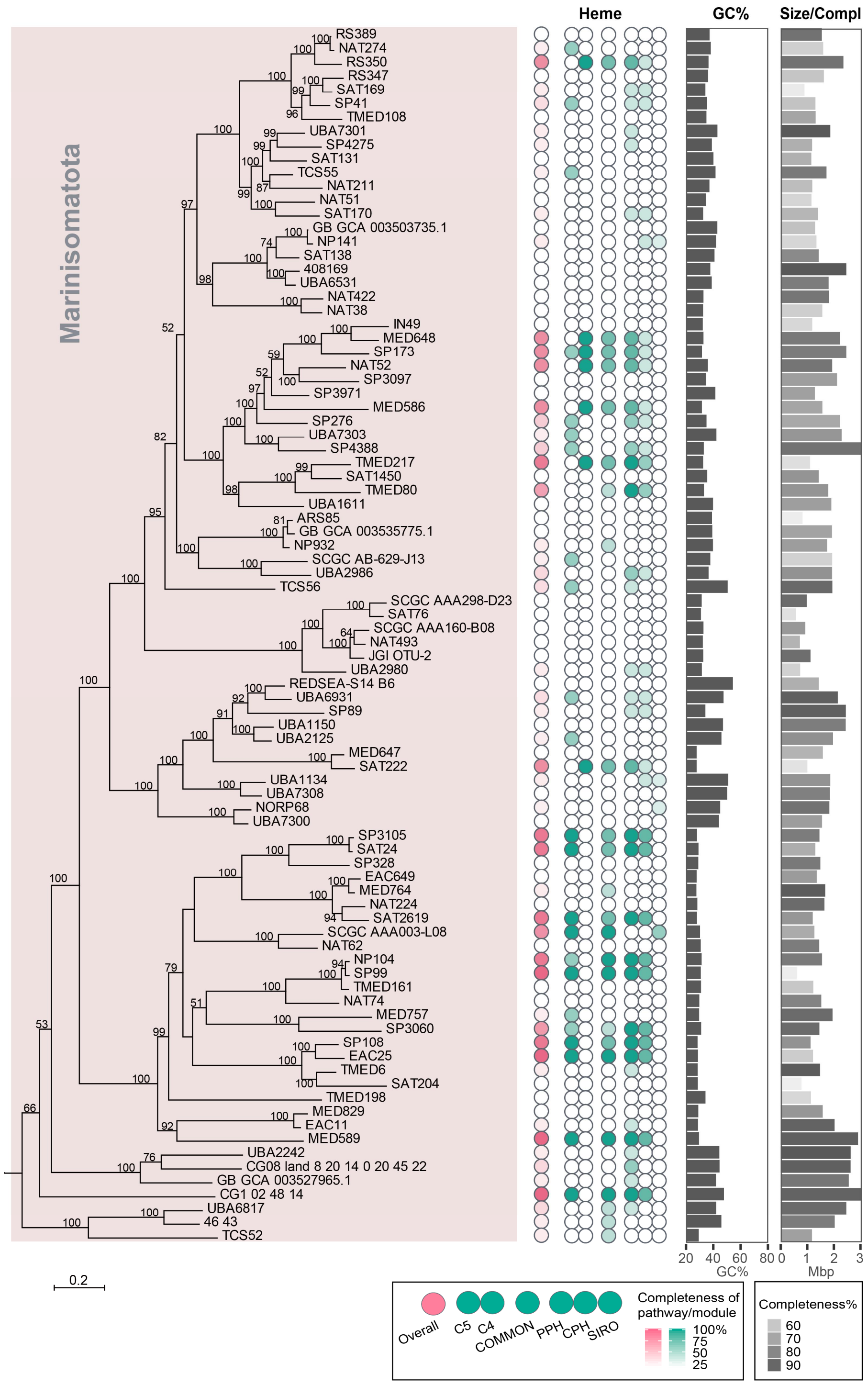


**Extended Data Fig. 7** Heme biosynthetic pathway completeness of representative genomes for species clusters belonging to the *Marinisomatota*. *Flavobacterium aquatile* LMG 4008 (not shown) was set as an outgroup in the tree on the left. Refer to the Extended Data Fig. 5 legend for a detailed explanation.

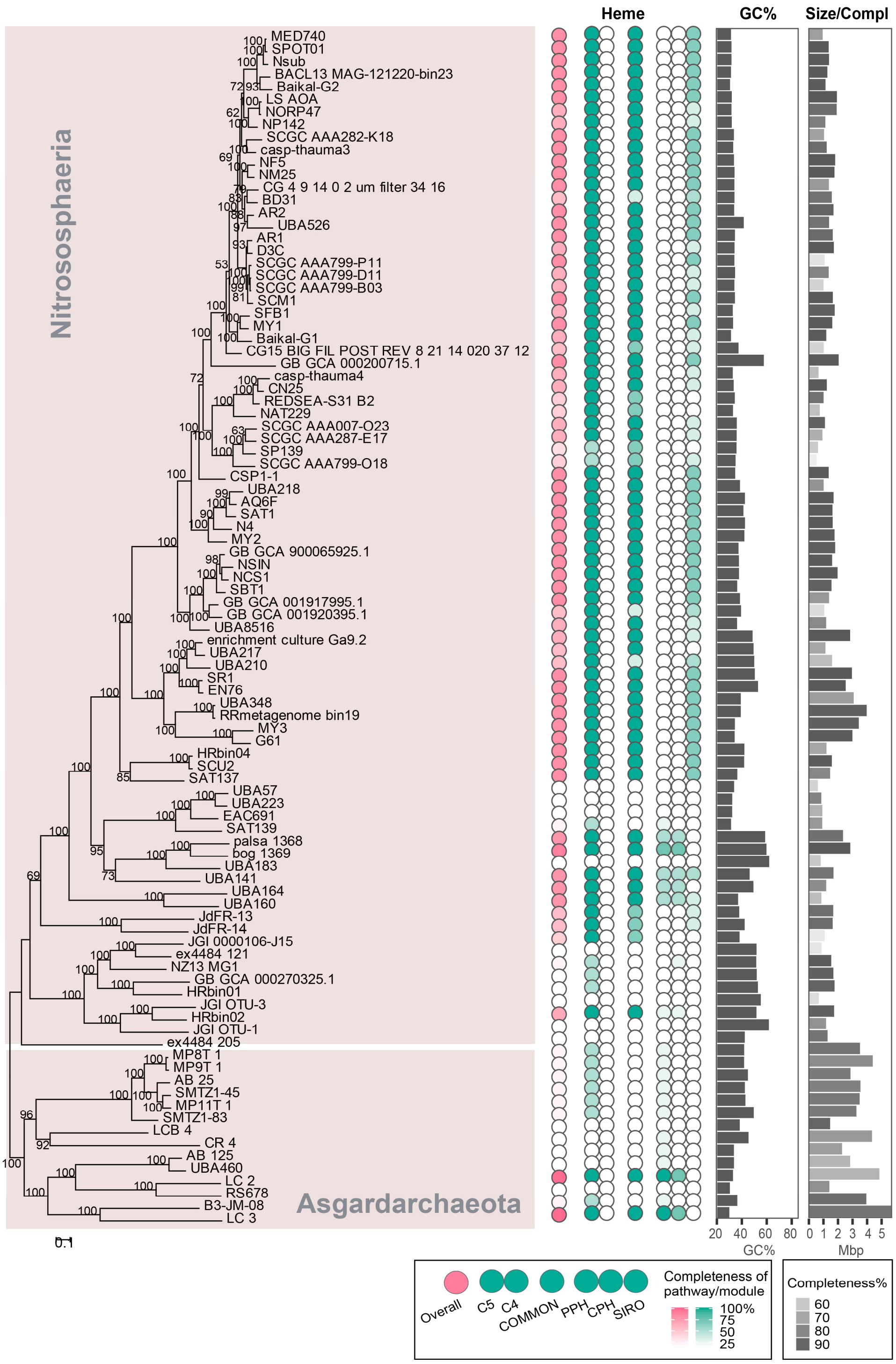


**Extended Data Fig. 8** Heme biosynthetic pathway completeness of representative genomes for species clusters belonging to the *Nitrososphaeria* and *Asgardarchaeota*. *Asgardarchaeota* was set as an outgroup in the tree on the left. Refer to the Extended Data Fig. 5 legend for a detailed explanation.

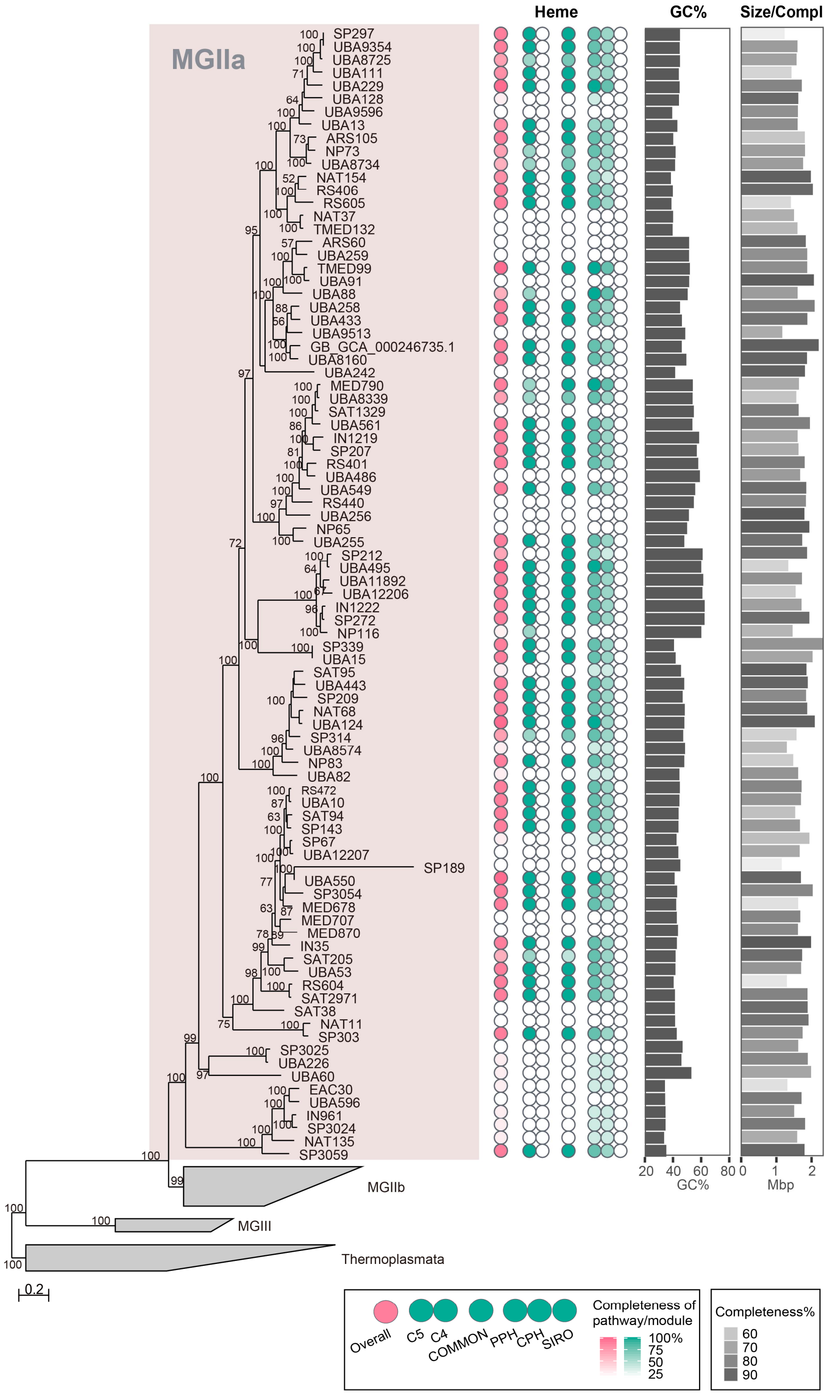


**Extended Data Fig. 9** Heme biosynthetic pathway completeness of representative genomes for species clusters belonging to MGIIa of *Archaea*. *Thermoplasmata* was set as an outgroup in the tree on the left. Refer to the Extended Data Fig. 5 legend for a detailed explanation.

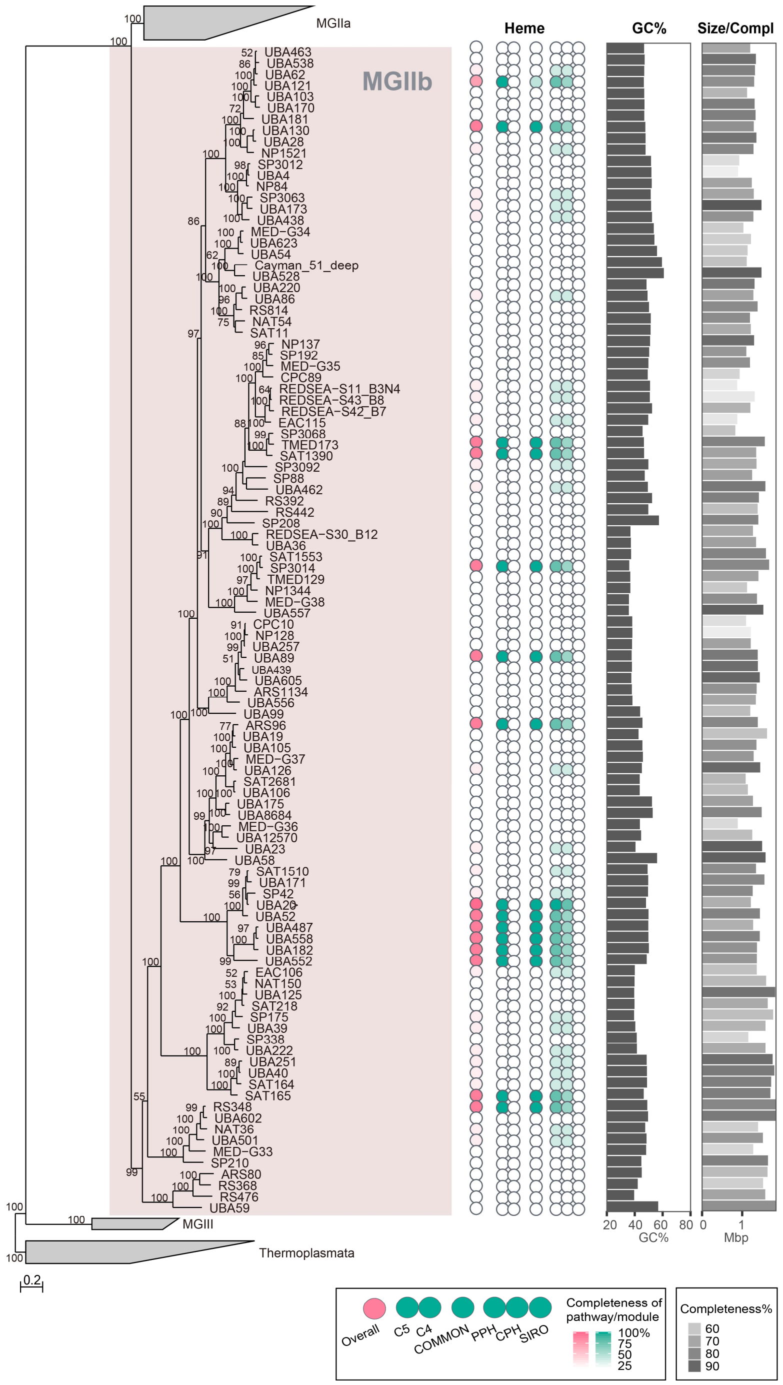


**Extended Data Fig. 10** Heme biosynthetic pathway completeness of representative genomes for species clusters belonging to MGIIb of *Archaea*. *Thermoplasmata* was set as an outgroup in the tree on the left. Refer to the Extended Data Fig. 5 legend for a detailed explanation.

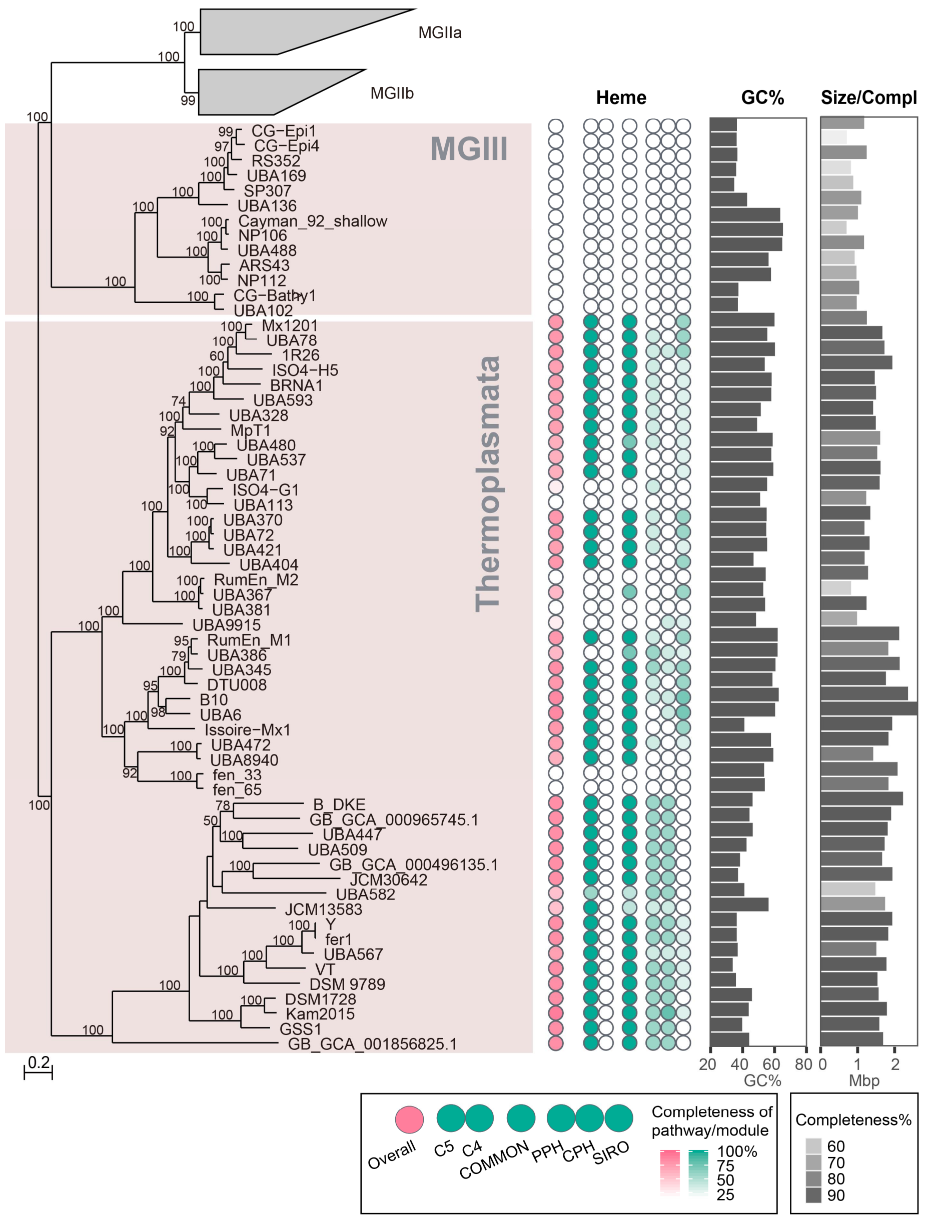


**Extended Data Fig. 11** Heme biosynthetic pathway completeness of representative genomes for species clusters belonging to MGIII and *Thermoplasmata* of *Archaea*. *Thermoplasmata* was set as an outgroup in the tree on the left. Refer to the Extended Data Fig. 5 legend for a detailed explanation.

**Extended Data Table 1** Composition of the medium used in this study

| Components | Compound(s) | Final concentration |
|---|---|---|
| Ammonium | $NH_4Cl$ | 10 µM |
| Phosphate | $KH_2PO_4$ | 10 µM |
| Trace metals | $FeCl_3 \cdot 6H_2O$ | 117 nM |
| | $MnCl_2 \cdot 4H_2O$ | 9 nM |
| | $ZnSO_4 \cdot 7H_2O$ | 800 pM |
| | $CoCl_2 \cdot 6H_2O$ | 500 pM |
| | $Na_2MoO_4 \cdot 2H_2O$ | 300 pM |
| | $Na_2SeO_3$ | 1 nM |
| | $NiCl_2 \cdot 6H_2O$ | 1 nM |
| Vitamin mixture | Thiamine·HCl | 59 nM |
| | Niacin | 81 nM |
| | Ca-Pantothenate | 84 nM |
| | Pyridoxine | 59 nM |
| | Biotin | 409 pM |
| | Folic acid | 453 pM |
| | Vitamin B12 | 70 pM |
| | Myo-inositol | 555 nM |
| | *p*-Aminobenzoic Acid | 7 nM |
| Carbon mixture | Pyruvate | 50 µM |
| | D-Glucose | 5 µM |
| | *N*-Acetyl-D-glucosamine | 5 µM |
| | D-Ribose | 5 µM |
| | Methyl alcohol | 5 µM |
| 20 proteinogenic amino acid mixture | Each amino acid | 100 nM, each |

**Extended Data Table 2** List of the microbial group names used in this study and their GTDB taxonomy

| Group name | Alternative | GTDB taxonomy |
|---|---|---|
| *Limnohabitans* | | d__Bacteria;p__Proteobacteria;c__Gammaproteobacteria;o__Burkholderiales;f__Burkholderiaceae;g__Limnohabitans |
| *Polynucleobacter* | | d__Bacteria;p__Proteobacteria;c__Gammaproteobacteria;o__Burkholderiales;f__Burkholderiaceae;g__Polynucleobacter |
| OM43 | | d__Bacteria;p__Proteobacteria;c__Gammaproteobacteria;o__Burkholderiales;f__Methylophilaceae;g__BACL14 |
| *Methylopumilus* | LD28 | d__Bacteria;p__Proteobacteria;c__Gammaproteobacteria;o__Burkholderiales;f__Methylophilaceae;g__Methylopumilus |
| *Halieaceae* | OM60/NOR5 | d__Bacteria;p__Proteobacteria;c__Gammaproteobacteria;o__Pseudomonadales;f__Halieaceae |
| *Spongiibacteraceae* | BD1-7 | d__Bacteria;p__Proteobacteria;c__Gammaproteobacteria;o__Pseudomonadales;f__Spongiibacteraceae |
| *Porticoccaceae* | SAR92 | d__Bacteria;p__Proteobacteria;c__Gammaproteobacteria;o__Pseudomonadales;f__Porticoccaceae |
| *Alteromonas* | | d__Bacteria;p__Proteobacteria;c__Gammaproteobacteria;o__Enterobacterales;f__Alteromonadaceae;g__Alteromonas |
| *Thioglobus* | SUP05 | d__Bacteria;p__Proteobacteria;c__Gammaproteobacteria;o__Thiomicrospirales;f__Thioglobaceae;g__Thioglobus |
| SAR86 | | d__Bacteria;p__Proteobacteria;c__Gammaproteobacteria;o__SAR86 |
| *Pelagibacterales* | SAR11, LD12 | d__Bacteria;p__Proteobacteria;c__Alphaproteobacteria;o__Pelagibacterales |
| HIMB59 | AEGEAN-169 | d__Bacteria;p__Proteobacteria;c__Alphaproteobacteria;o__HIMB59 |
| *Puniceispirillales* | SAR116 | d__Bacteria;p__Proteobacteria;c__Alphaproteobacteria;o__Puniceispirillales |
| *Rhodobacteraceae* | | d__Bacteria;p__Proteobacteria;c__Alphaproteobacteria;o__Rhodobacterales;f__Rhodobacteraceae |
| SAR324 | Marine group B | d__Bacteria;p__SAR324 |
| *Flavobacteriaceae* | | d__Bacteria;p__Bacteroidota;c__Bacteroidia;o__Flavobacteriales;f__Flavobacteriaceae |
| *Marinimicrobia* | SAR406, Marine group A | d__Bacteria;p__Marinisomatota |
| OM1 | | d__Bacteria;p__Actinobacteriota;c__Acidimicrobiia;o__TMED189 |
| acIV | | d__Bacteria;p__Actinobacteriota;c__Acidimicrobiia;o__Microtrichales;f__Ilumatobacteraceae;g__UBA3006<br>d__Bacteria;p__Actinobacteriota;c__Acidimicrobiia;o__Microtrichales;f__Ilumatobacteraceae;g__BACL27<br>d__Bacteria;p__Actinobacteriota;c__Acidimicrobiia;o__Microtrichales;f__Ilumatobacteraceae;g__UBA2093<br>d__Bacteria;p__Actinobacteriota;c__Acidimicrobiia;o__Microtrichales;f__Ilumatobacteraceae;g__Casp-actino8 |
| *Rhodoluna* | | d__Bacteria;p__Actinobacteriota;c__Actinobacteria;o__Actinomycetales;f__Microbacteriaceae;g__Rhodoluna |
| acI | | d__Bacteria;p__Actinobacteriota;c__Actinobacteria;o__Nanopelagicales;f__Nanopelagicaceae |
| SAR202 | | d__Bacteria;p__Chloroflexota;c__Dehalococcoidia;o__UBA1151<br>d__Bacteria;p__Chloroflexota;c__Dehalococcoidia;o__SAR202<br>d__Bacteria;p__Chloroflexota;c__Dehalococcoidia;o__UBA3495<br>d__Bacteria;p__Chloroflexota;c__Dehalococcoidia;o__UBA2985<br>d__Bacteria;p__Chloroflexota;c__Dehalococcoidia;o__GCA-2717565<br>d__Bacteria;p__Chloroflexota;c__Dehalococcoidia;o__UBA2963<br>d__Bacteria;p__Chloroflexota;c__Dehalococcoidia;o__UBA6952<br>d__Bacteria;p__Chloroflexota;c__Dehalococcoidia;o__UBA1127<br>d__Bacteria;p__Chloroflexota;c__Dehalococcoidia;o__SAR202-VII-2 |
| CL500-11 | | d__Bacteria;p__Chloroflexota;c__Anaerolineae;o__Anaerolineales;f__UBA11657 |
| *Patescibacteria* | Candidate phyla radiation | p__Patescibacteria |
| *Asgardarchaeota* | Asgard archaea | p__Asgardarchaeota |

| | | |
|---|---|---|
| Marine Group IIa | | d__Archaea;p__Thermoplasmatota;c__Poseidoniia;o__Poseidoniales;f__Poseidoniaceae |
| Marine Group IIb | | d__Archaea;p__Thermoplasmatota;c__Poseidoniia;o__Poseidoniales;f__Thalassoarchaeaceae |
| Marine Group III | | d__Archaea;p__Thermoplasmatota;c__Poseidoniia;o__MGIII |
| Thaumarchaeota | Marine Group I | d__Archaea;p__Crenarchaeota;c__Nitrososphaeria |

**Extended Data References**

1. Haase, I. et al. Enzymes from the haloacid dehalogenase (HAD) superfamily catalyse the elusive dephosphorylation step of riboflavin biosynthesis. *Chembiochem* **14**, 2272-2275 (2013).

2. Sarge, S. et al. Catalysis of an essential step in vitamin B2 biosynthesis by a consortium of broad spectrum hydrolases. *Chembiochem* **16**, 2466-2469 (2015).
